# Relating Polar Bears Killed, Human Presence, and Ice Conditions in Svalbard 1987 – 2019

**DOI:** 10.1101/2023.03.17.533082

**Authors:** D. Vongraven, S.C. Amstrup, T.L. McDonald, J. Mitchell, N.G. Yoccoz

**Affiliations:** Department of Arctic and Marine Biology, UiT Arctic University of Norway, N-9037 Tromsø, Norway; Norwegian Polar Institute, Fram Center, N-9296 Tromsø, Norway; Polar Bears International, 810 N. Wallace, Suite E, Bozeman, MT 59715-3020, USA; McDonald Data Sciences, 1529 Rainbow Ave, Laramie, WY 82070, USA; Western Ecosystems Technology, Inc., 415 W. 17^th^ Street, Suite 200, Cheyenne, WY 82001, USA

## Abstract

Conflicts between humans and polar bears have been predicted to increase as polar bear prime habitat, sea ice, is decreasing. In Svalbard, a strong protection and strict control schemes have secured near complete records of bears killed and found dead since 1987. We analyzed the trend in the number of kills and related this to human visitation to the island. We found a slight decrease in the number of kills in the period 1987-2019, and a decrease in per capita number of kills when monthly kills were compared to the monthly number of visitors disembarking in the main settlement. We then used a discrete choice resource selection model to assess whether polar bear kill events are related to attributes of the kill sites and environmental conditions at the time. We divided Svalbard in four sectors, North, East, South, and West, and monthly average ice cover was calculated in 25-km rings around Svalbard, rings that were further delineated by the four sectors. We found that the odds of a kill was greater along the shoreline, and that the odds would be reduced by 50% when moving only 900 m from the shoreline when all sectors were included. Distance from other covariates like settlements, trapper’s cabins, and landing sites for tourists did for the most part not have a significant impact on the odds of a kill. Sectorwise, ice cover had no significant impact on the odds for a kill. The decreasing trend in kills of polar bears might partly be explained by the success of strict protection and management regimes of Svalbard wilderness.

## INTRODUCTION

On a global scale, habitat loss is one of the most critical threats to persistence of mammal populations (Schipper et al., 2008). This is especially apparent for species dependent on Arctic sea ice habitat which has shown significant spatial and temporal declines with global climate warming (Stroeve et al., 2014; Laidre et al., 2015; Stern and Laidre, 2016; National Snow and Ice Data Center, 2019). In the case of polar bears (*Ursus maritimus*), large ice dependent predators, sea ice decline will have a direct effect on the extent and characteristics of preferred habitat (Durner et al., 2009; Laidre et al., 2018; Lone et al., 2018) the availability of preferred prey (Pagano et al., 2018), and the amount of time they are able to stay on sea ice to find prey (Atwood et al., 2016), all which could lead to increased contact with humans and thus increased number of conflicts with humans (Stirling and Derocher, 2012; Atwood and Wilder, 2021; Rode et al., 2022; Abrahms et al., 2023).

Polar bears depend on sea ice for fundamental aspects of their life history, breeding, accessing preferred denning areas, and most importantly foraging (Stirling, 1974; Amstrup et al., 2008; Derocher et al., 2011). Arctic Ocean September sea ice extent has declined by 81,200 km^2^/year since 1979, i.e. a 12.7% (+/- 2.0%) decline per decade (Meier and Stroeve, 2022). In the Barents Sea, the ice-free period between spring melt and fall freeze-up, has increased 34 days per decade since 1979 (Stern and Laidre, 2016). Historically, sea ice advanced from the northeast in November and engulfed the Svalbard Archipelago through August (Divine and Dick, 2006). However, in the period 1979-2022 there was an observed negative trend of 10.1% per decade in March sea ice extent, and a 19.7% per decade reduction in September sea ice extent (https://cryo.met.no/en/sea-ice-index-bar), leading to little or no sea ice surrounding Svalbard for most of the year. Also, most of 29 fjord regions in Svalbard show decadal decrease of ice cover in winter months of 10-40% in the two decades prior to 2016, whilst most showed decadal slight increase in ice cover in the two decades prior to 1998 (Dahlke et al., 2020).

Similar patterns have been documented in other Arctic areas. In Alaska, for example, sea ice retreat forced bears to swim long distances between shore and sea ice in summer, in some instances with fatal consequences (Monnett and Gleason, 2006; Durner et al., 2011; Pagano et al., 2012), and spend more time on land in fall (Rode et al., 2015). In Hudson Bay, where a longer ice-free period has led to decline in body condition for adult males and females in both the Western and Southern Hudson Bay subpopulations (Stirling et al., 1999; Obbard et al., 2018), cub survival and population size in Western Hudson Bay have declined (Regehr et al., 2007; Lunn et al., 2016). Energy budget modeling demonstrates limits to how long polar bears can be food deprived before survival is affected, ranging from 117 days in females with cubs till 255 days in solitary females that start fasting in average body condition (Molnár et al., 2020). Severe global declines in polar bear numbers are projected if global warming induced sea ice loss continues (Amstrup et al., 2010; Regehr et al., 2016; Molnár et al., 2020).

Growing human populations, industrial activity, and tourism across the polar bear’s range exacerbate potential for conflict (Anonymous, 2009; Larsen and Fondahl, 2015; Atwood et al., 2017). The number of people arriving in Svalbard to visit the Norwegian Arctic has been growing for decades; visitation to Svalbard has increased nearly 8-fold since the 1990s (Bystrowska, 2019; Norwegian Polar Institute, 2022). More people visiting bear habitat coupled with more bears on land due to decreasing sea ice cover raise the potential for both an increase in human-bear conflict locally, e.g. Western Hudson Bay (Towns, 2006; Towns et al., 2009), and for the severity of such conflicts globally (Can et al., 2014).

Here, we examine kills of problem bears in Svalbard to determine whether ongoing environmental changes, including the sea ice concentration at the time of the kill, may have influenced their frequency or distribution. We used a discrete choice resource selection model (Cooper and Millspaugh, 1999; Manly et al., 2002; McDonald et al., 2006) to assess how polar bear kill events are related to attributes of the kill sites and environmental conditions at the time.

## MATERIAL AND METHODS

### Data on killed bears

Strict protection and control schemes assure records were kept for all killed or found dead polar bears. We initially obtained records of polar bears killed or found dead from the Governor of Svalbard for the period August 1987 to August 2016. We later obtained one more record of killed bear in July 2018, and there were no killed or dead bears on record for 2019. Data always included date and location. In most cases, records also contained sex and age of the bear, whether it was previously tagged for scientific purposes, a narrative giving details about the incident, and the people involved. In most cases prior to 2005 the given coordinates did not have GPS accuracy. In all cases the narratives were checked in order to set the coordinates for the recorded incident as accurate as possible.

We based our analysis on the initial set of records covering August 1987 to August 2016 that contained 113 bear deaths. We removed 18 records from this data set. We discarded nine records that did not have adequate information or for which we suspected data entry errors. One record was duplicated, the only record where a data entry error was clearly detectable. One bear with satellite collar that was found dead and cause of death could not be determined. We removed records of two bears that were euthanized due to severe injuries. We removed one record of a dead cub from a killed mother-cub pair. We removed four euthanized cubs whose mothers were killed as DLP, but we retained the mother’s DLP records. These initial exclusions left 95 bears in the statistical analysis of kill locations (Figure 1).

**Figure 1.**
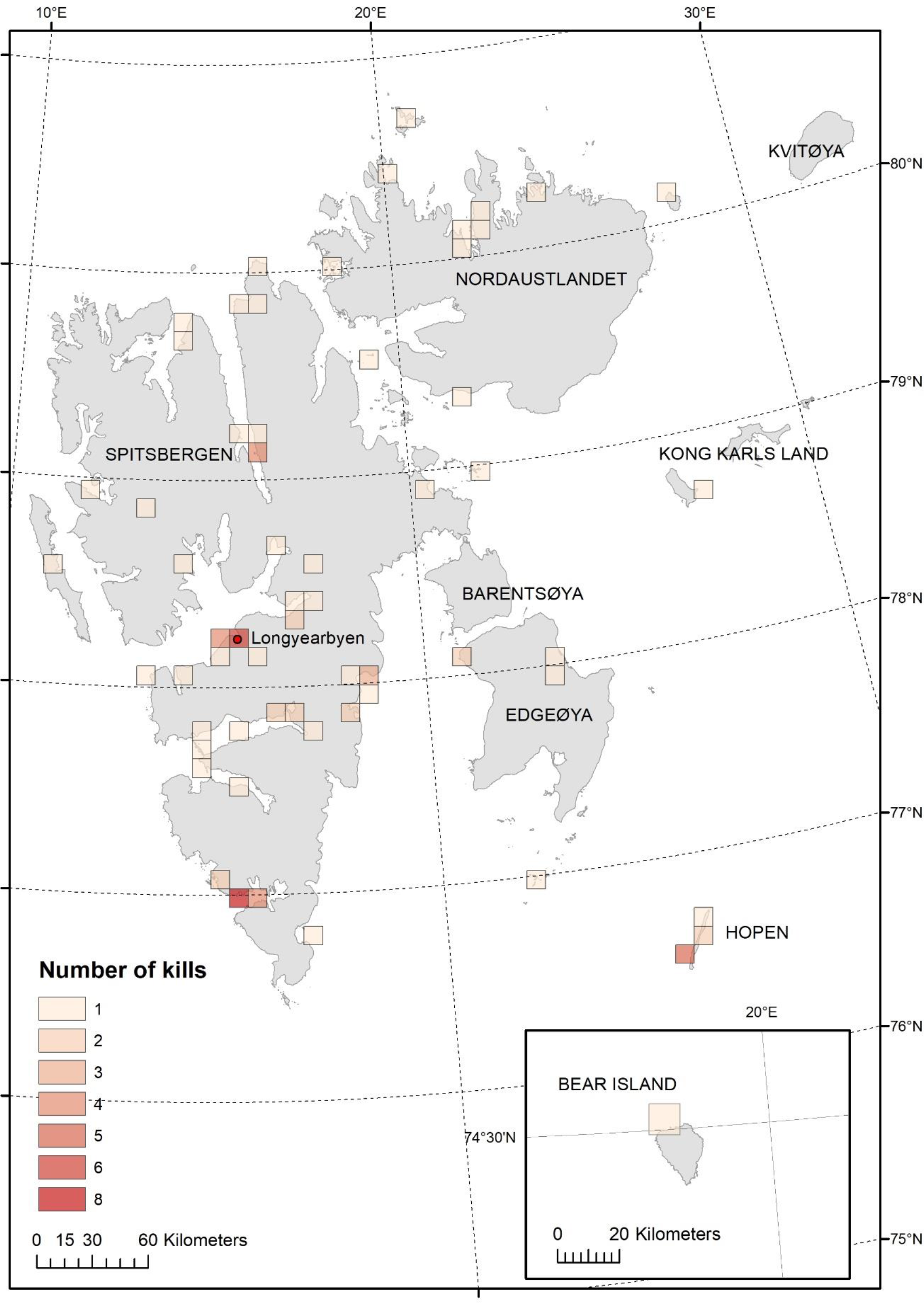
Killed polar bears and polar bears found dead in Svalbard 1987-2016, map showing 95 locations aggregated over at 10 km grid

Eighty-one of the 95 records, covering the period August 1987 – August 2016 were classified as problem bears, or bears legally killed in defence of life and property (also called DLP-bears, Supplement S1). A problem bear is a bear that obtained anthropogenic food, damaged property, acted aggressively towards humans, or was judged to be negatively affected by human activities (Hopkins et al., 2010; Wilder et al., 2017). Of the 14 bears classified as “non-DLP”, six were natural mortalities and eight were sick or injured bears killed by the government for ethical reasons (Supplement S1). The two euthanized bears removed from the original 113 bear dataset should have been included in the “non-DLP” data, however, when this error was discovered analyses were already finalized, and we concluded that these records would have negligible influence on the results.

Analyses was first done separately for dataset “DLP” (81 records) and dataset “ALL” (95 records), but since the results were similar we only describe results from the data set “ALL”.

In late 2021, we obtained data for 2017-2019, and there were one more record of a kill that occurred in 2018, none in 2017 or 2019. We did not include this kill in our statistical modelling of kill location, but we did include it in other analyses and tallies of the number of kills. Hence, the dataset used to report total number of kills consisted of 96 bears, covering the period August 1987 and through 2019.

Of the six data sets on tourism in Svalbard found available (Table S2), we utilized the data set showing monthly totals of the number of people arriving in Longyearbyen from 1995 to present (Figure S3). These data were compared to the monthly numbers of kills for all sectors combined.

### Analyses of the numbers of killed bears

To examine for trends in annual counts of killed bears we analysed the yearly counts using a generalized linear model (GLM) with a Poisson distribution and a log link. We used quasi-likelihood methods to account for overdispersion in the data (assessed using the residual deviance; McCullagh and Nelder (1989)), and checked if residuals of the models were autocorrelated using the acf() function in R. To examine changes among single months and whether there were changes in single months over years we used a GLM on the month by year counts and the aggregated monthly counts.

We assessed relationships between number of bears killed and number of tourists, with a log-linear model, assuming a Poisson distribution for the response and a log link. We split 1995-2019 in three equal length periods (1995-2002, 2003-10, and 2011-2019) to assess temporal changes in the relationship between numbers of killed bears and numbers of tourists. We tested for an interaction between period and number of tourists, and then for the additive effect of period and number of tourists (each effect fitted last). Overdispersion of the most complex model (with the interaction period*number of tourists) was assessed using the residual deviance (McCullagh and Nelder, 1989). Standardized residuals were checked for trends and constant variance. These analyses were done in R (R Core Team, 2022).

### Analyses of kill locations and covariates

We applied discrete-choice analysis to examine variables hypothesized to influence the odds of a kill event in a specific location (Cooper and Millspaugh, 1999). Discrete-choice models compare characteristics of kill locations to characteristics of random locations and are a variant of conditional logit models (Allison, 1999). Our discrete choice model related the odds of a kill (odds = probability of kill divided by probability of no kill) to changes in the explanatory variables. In particular, we were interested in whether changes in offshore ice concentrations were correlated with odds of a kill after controlling for other sources of variation such as distance to shore and settlements.

The underlying statistical likelihood of our discrete choice model is the same as that utilized in stratified Cox proportional hazards modelling of continuous-time survival rates, where stratum are composed of one case (here, kill) and several controls (here, random location) (Therneau and Grambsch, 2000). Thus, we used statistical software procedures capable of estimating stratified Cox proportional-hazards models to estimate our discrete-choice models (McDonald et al., 2006; Therneau, 2017).

For the discrete choice analyses, we required random locations to compare against every kill location. We generated 50 unique random locations for each kill event on the landmass of Svalbard (map scale 1 : 250,000; map data from Norwegian Polar Institute, 2014). Random points were chosen using ArcMap software, with random points generated from a uniform distribution over triangles that partitioned area polygons (ESRI, ArcMap 10.3). All random points were on land even though 11 of the 95 kills were slightly (<600 m) offshore and three were farther (1304, 2868 and 5559 m offshore). All 14 offshore kill sites were within narrow fjords of the Svalbard archipelago. In total, the DLP dataset used for modelling contained (50 random + 1 kill) x (81 sites) = 4,131 points, while the ALL dataset contained (50 random + 1 kill) x (95 sites) = 4,845 points. All random points had the same time stamp as their assigned so that temporal characteristics (e.g., tourist landings) could be correctly assigned.

We derived five distance metrics from the locations of random points and kills. For each of the random and kill-site points we measured the shortest straight-line distance to sea (SEA), distance to the nearest tourist landing location in use during 1996-2016 (LAND124; Figure 2A), distance to the nearest of the 21 large landing sites that welcomed >10,000 tourists during the same period (LAND21; Figure 2A), distance to the nearest of seven permanent settlements (SET; Figure 2B), and the distance to the nearest of five trapper’s cabins that were active during the period (TRAP; Figure 2B). We treated all 124 landing sites as active during the period 1987-2016 even though the data on landings were available only after 1995 and some were intermittently inactive thereafter. When inactive, zero landings were recorded at landing sites. Similarly, we assumed all settlements were equivalent even though few people resided in some settlements while Longyearbyen, the largest settlement, had more than 2,100 residents annually.

**Figure 2.**
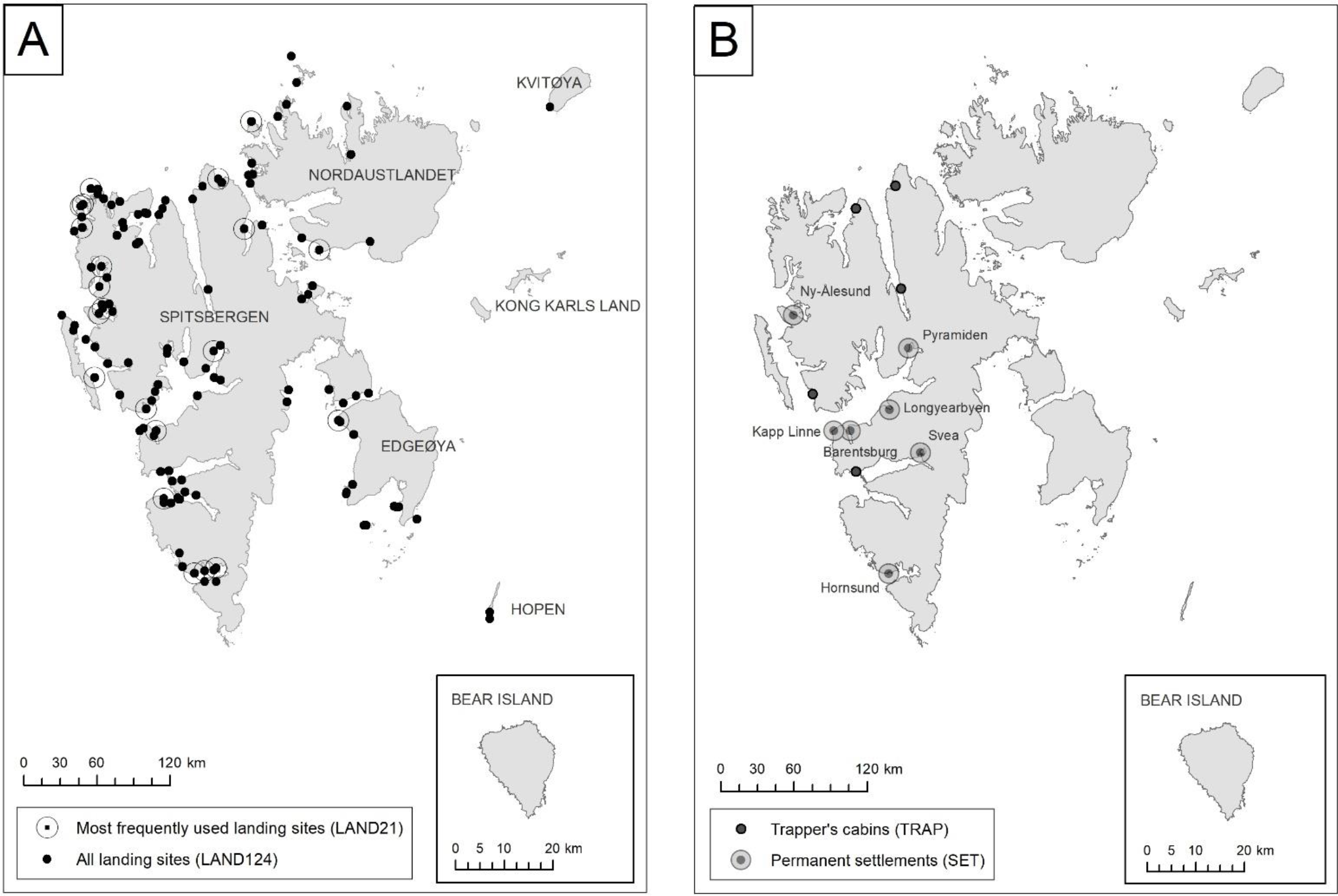
Locations of human activity used to calculate distance covariates. Large caps annotate land masses/islands, small caps annotates settlements.

Pearson correlations amongst the set of all kill and random sites demonstrated that LAND124, LAND21, SET, and TRAP were moderately inter-correlated (0.41 ≤ r ≤ 0.76). To prevent multicollinearity and model parameter instability of discrete-choice models, we included up to two distance metrics, including only one from the set of four correlated distance metrics, and potentially the uncorrelated distance to sea metric (SEA).

To investigate regional differences, we assigned every kill and random point to one of four geographic sectors covering Svalbard and the surrounding sea. We aligned our sectors with the cardinal directions west (W), south (S), east (E), and north (N) because ice flow and coverage were generally similar within and dissimilar among sectors (Figure 3A). Sectors varied in size and the W sector encompassed the main settlement Longyearbyen.

**Figure 3.**
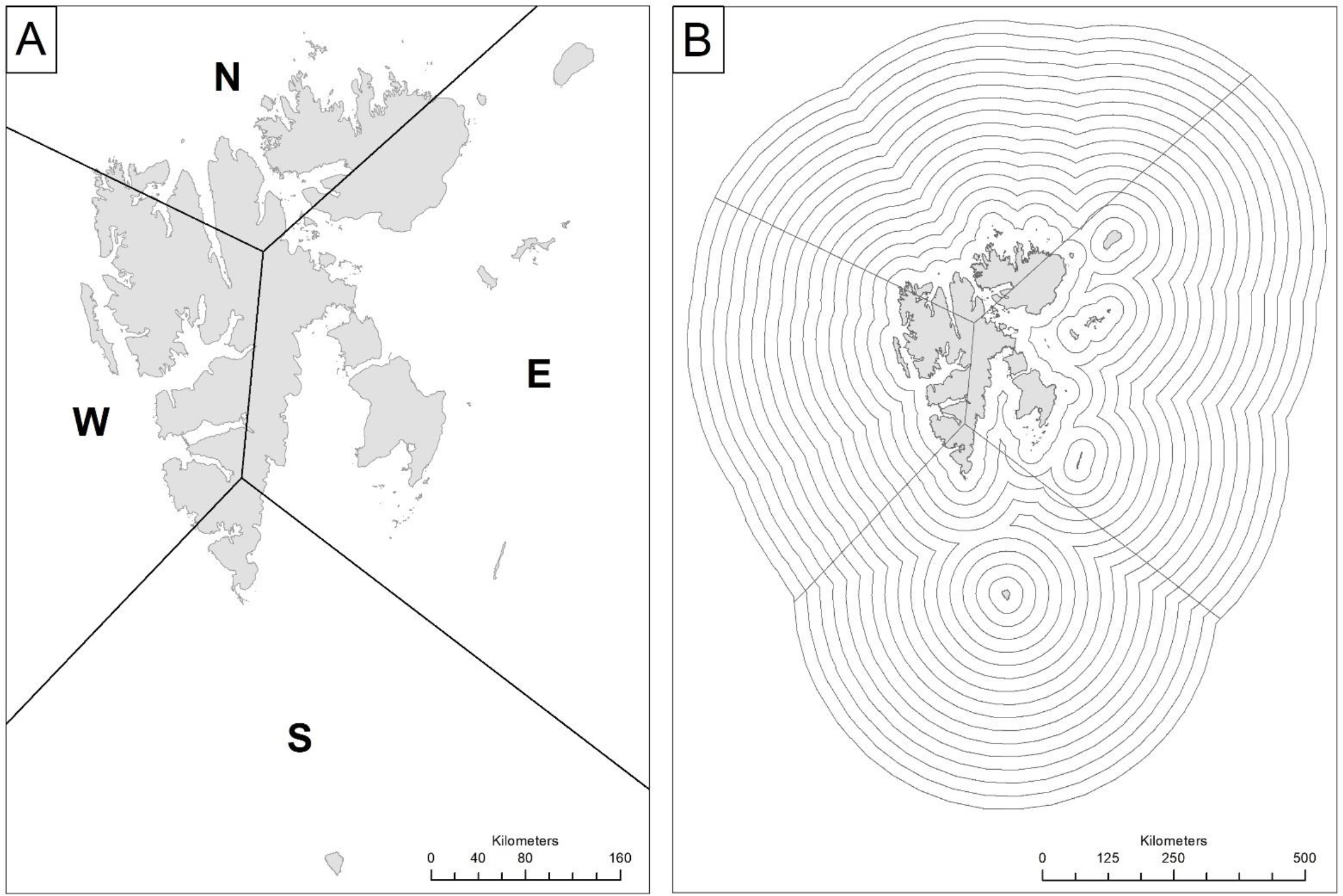
Panel A shows the arbitrarily chosen sectors and panel B the 25-km wide bands used for aggregation of monthly sea ice concentration 1987-2016.

To determine whether offshore sea ice concentration influenced probability of a bear kill, we measured ice concentration in 25-km wide radial bands surrounding the Svalbard shoreline out to a distance of 400 km (Figure 3B). We obtained daily measurements of sea ice concentration out to 400 km from the National Snow and Ice Data Center (NSIDC, Boulder, Colorado, USA). We calculated monthly average sea ice concentration separately for each 25-km band in each sector using data from November 1978 through December 2016, thus ending up with 16 ice cover covariates for each data set, whether being for each separate sector or for all sectors combined.

To investigate whether the odds of any particular depended on ice cover in months prior to the kill (ice lag), we introduced lagged months of ice cover as a covariate. Monthly lag periods ranged from zero to nine months. A lag of zero months means ice coverage data from the month in which the bear was killed was used as predictor, while a lag of nine months means the ice-coverage data used was from nine months prior to the bear-kill date. Note that nine months safely encompasses any one year’s maximum fall freeze-up and spring melt. We considered all 10 monthly lagged models, from the zero-month lag model to the 9-month lag model.

Evaluation of at most one of the 16 different ice-cover covariates; either zero, one, or two distance metrics, induced a total number of up to 169 possible models, although sparse data occasionally prevented the evaluation of some of these possibilities. Assuming convergence, we used Akaike’s Information Criteria (AIC) to determine which models were better predictive models than others (Anderson and Burnham, 2011).

Five different data subsets were considered for model evaluation. We fitted one model to a dataset containing records from all sectors. We fitted separate models to data from each sector to allow sector-specific covariates to manifest when appropriate. Sector-specific models were subsetted to only include a kill originating in the sector of interest. Once identified, the 50 replicates tied to those sector-specific bears were also included. In this way, the four separate sector-specific models partitioned the available data. We then fitted one model to data from all sectors combined. AIC identified best models on each of the five different datasets independently.

We obtained odds ratios by exponentiating estimated model coefficients. Odds ratios measured the direction and magnitude of changes in the probability of a kill occurring in a pixel when the value of a covariate changes by one unit (we used the original scale for all covariates). For example, an odds ratio of 0.5 for distance from sea (covariate SEA) meant that the odds (probability of a kill divided by probability of no kill) decreased by half for each one kilometer increase in distance to the sea. Conversely, the odds ratio’s reciprocal, 1/0.5 = 2, implies that a pixel three kilometer from the sea will have twice the odds of containing a kill location as a pixel two kilometer from the sea (assuming all else is the same) because these two locations are one kilometer apart. We considered odds ratios statistically significant if their 95 % confidence interval did not include 1. For reporting, we calculated the theoretical distance a bear would be required to move away from three types of locations (shoreline, landing site, and settlement) to reduce or augment their odds for being killed by 50%. We also calculated the change in ice cover in a given ring for a given lag that would change the odds for a kill by 50%.

## RESULTS

### Number of killed bears

Annual polar bear kills during the period 1987-2019 ranged from 0-10. Evidence for an apparently declining trend in annual kills was weak after accounting for overdispersion (b_year_=-0.026 [95% CI: -0.054; 0.002]; F_1,26_=3.27, P= 0.08, Figure 4, panel A). Variation among months, however, was great (ΔDeviance=59.15, df=11, P<0.001) with peaks in bear kills in April, July and August, and low numbers in October and November (Figure 4, panel B). Kills in individual months showed no strong evidence for long-term trend (all P>0.08). August, the month with most kills, reflected the same negative trend as yearly data (b_year_=-0.036 [95% CI: -0.081; 0.004]).

**Figure 4.**
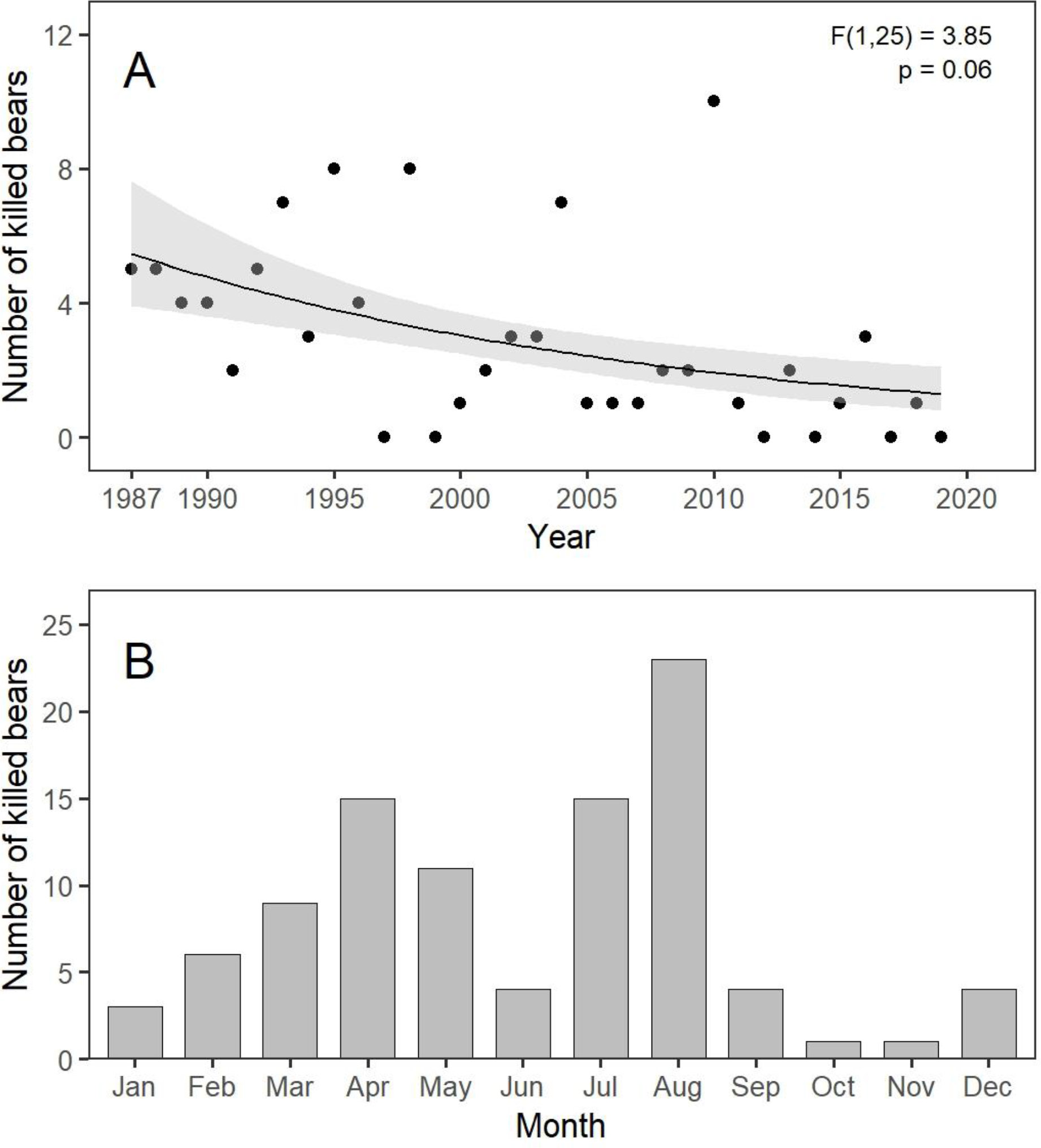
Polar bear kills in Svalbard 1986-2019 aggregated on a yearly (A) and monthly (B) scale. Annual best-fit trend line in upper panel, with 95% CI (grey area), obtained via a Poisson Generalized Linear Model with a quasilikelihood variance adjustment.

There was a positive relationship between the number of bears killed and the number of people arriving in Longyearbyen (Figure 5). The model with the interaction Tourists*Period fitted the data (residual deviance = 28.1, df = 30, p=0.57). There was no evidence for an interaction Tourists*Period (Δdeviance=1.32, df=2, p=0.52), meaning that the slope of the relationship log(mean number of bears killed) – number of tourists could be assumed to have been the same among periods. There was strong evidence when fitting the effect last for an additive effect of Period (Δdeviance=40.2, df=2, p<0.0001) and Number of Tourists (Δdeviance=29.2, df=1, p<0.0001). The period effect showed a strong decrease (Δ(Period2-Period1)=-0.74, SE=0.34, Δ(Period3-Period1)=-3.75, SE=0.78, i.e. number of bears killed in period 2 was ca halved compared to period 1 for the same number of tourists, and 42 times less in period 3 compared to period 1). Therefore, the overall increase in the number of people coming to Svalbard was compensated by a decreasing number of bears killed for a given number of people, leading to no systematic increase in the total number of bears killed over time.

**Figure 5.**
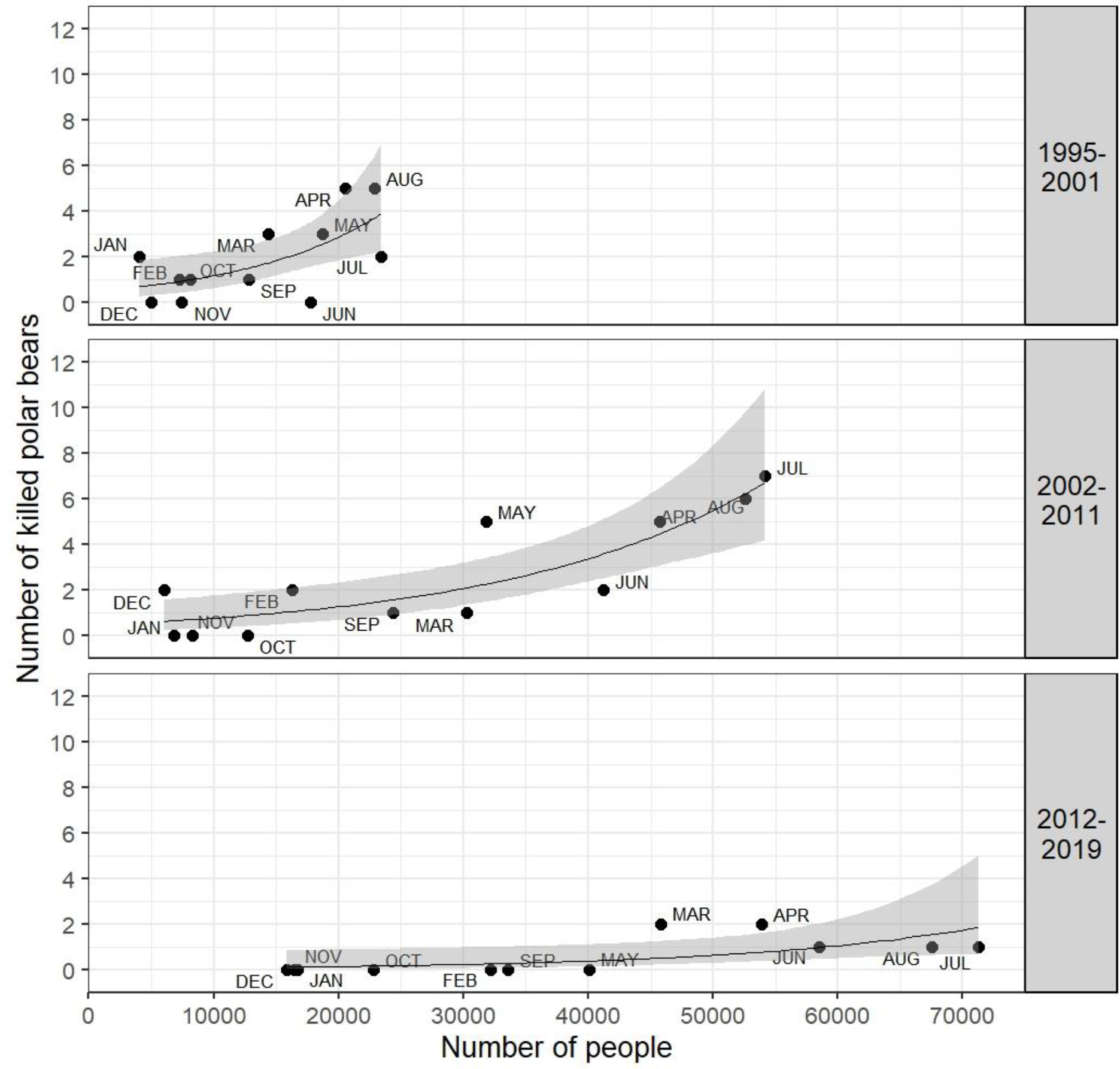
The relation between the number of tourists arriving in Longyearbyen and the number of bears killed in Svalbard, in three periods in the period 1987-2019. Annual best-fit trend lines, with 95% CI (grey area), obtained via a Poisson Generalized Linear Model with a quasilikelihood variance adjustment.

### Locations of killed bears

Figure 6 shows the spatial and temporal distributions of kills (number of kills between sectors is provided in Table S4). There were consistently more kills in August in all sectors, and there was a small peak centred on April when all sectors were combined. There were similar temporal patterns in all sectors, with a slight exception for the West-sector, where there was a higher number of kills than in the other sectors, and where the temporal distribution of kills was more uniform across months.

**Figure 6.**
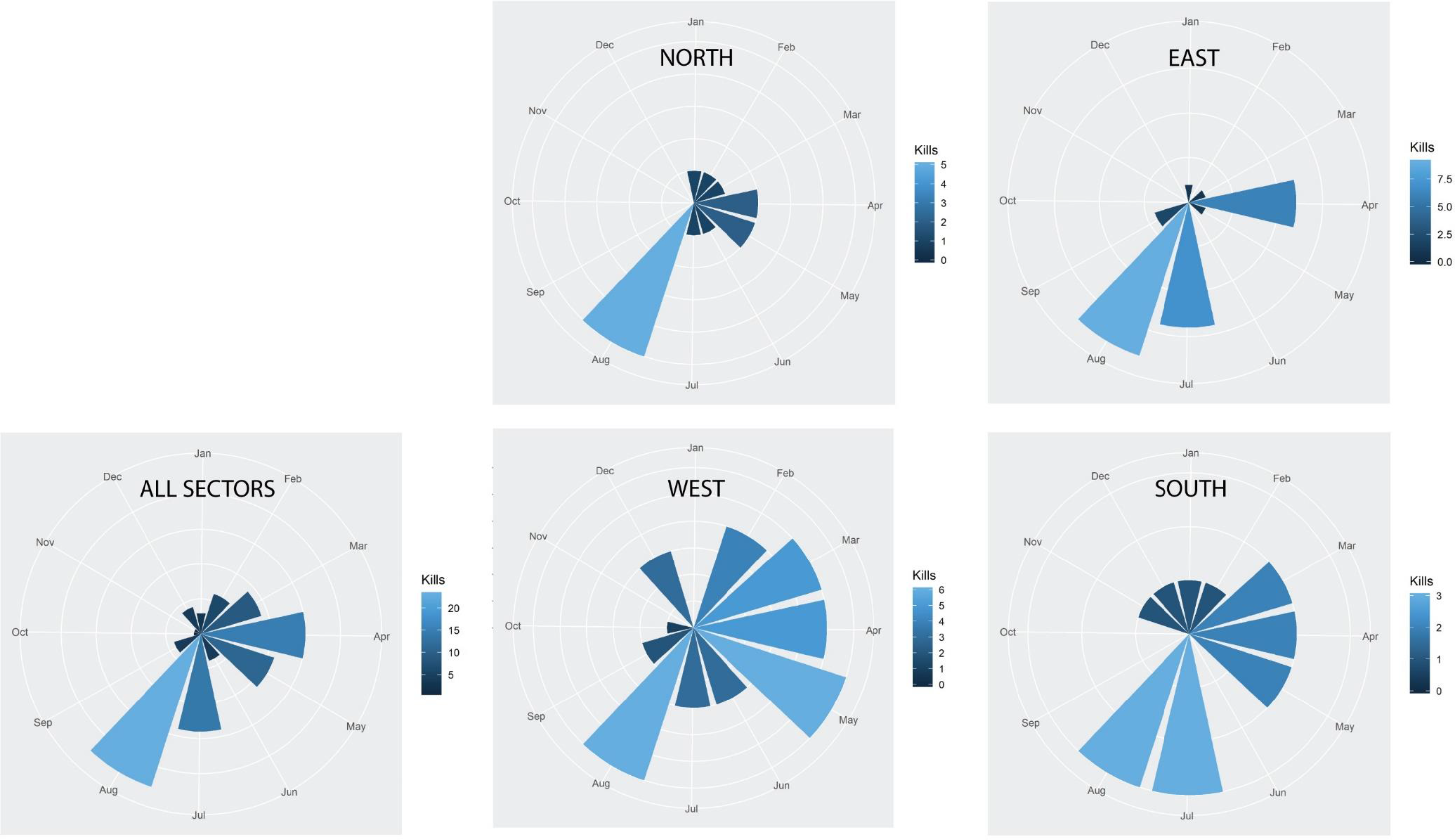
The timing of polar bear kills over All Sectors combined and per sector.

Tables 1A and 1B present the calculated additional average distance a theoretical bear would be required to move away from the shore line, a landing site, or a settlement in order to reduce their odds of being killed by 50% (Table S5 displays the corresponding odds ratios and confidence intervals from all the top model runs for all sectors and all the sectors combined, i.e. Sector All). Table 1C presents the same for change in ice cover resulting in a 50% reduction in odds of a kill.

**Table 1.**
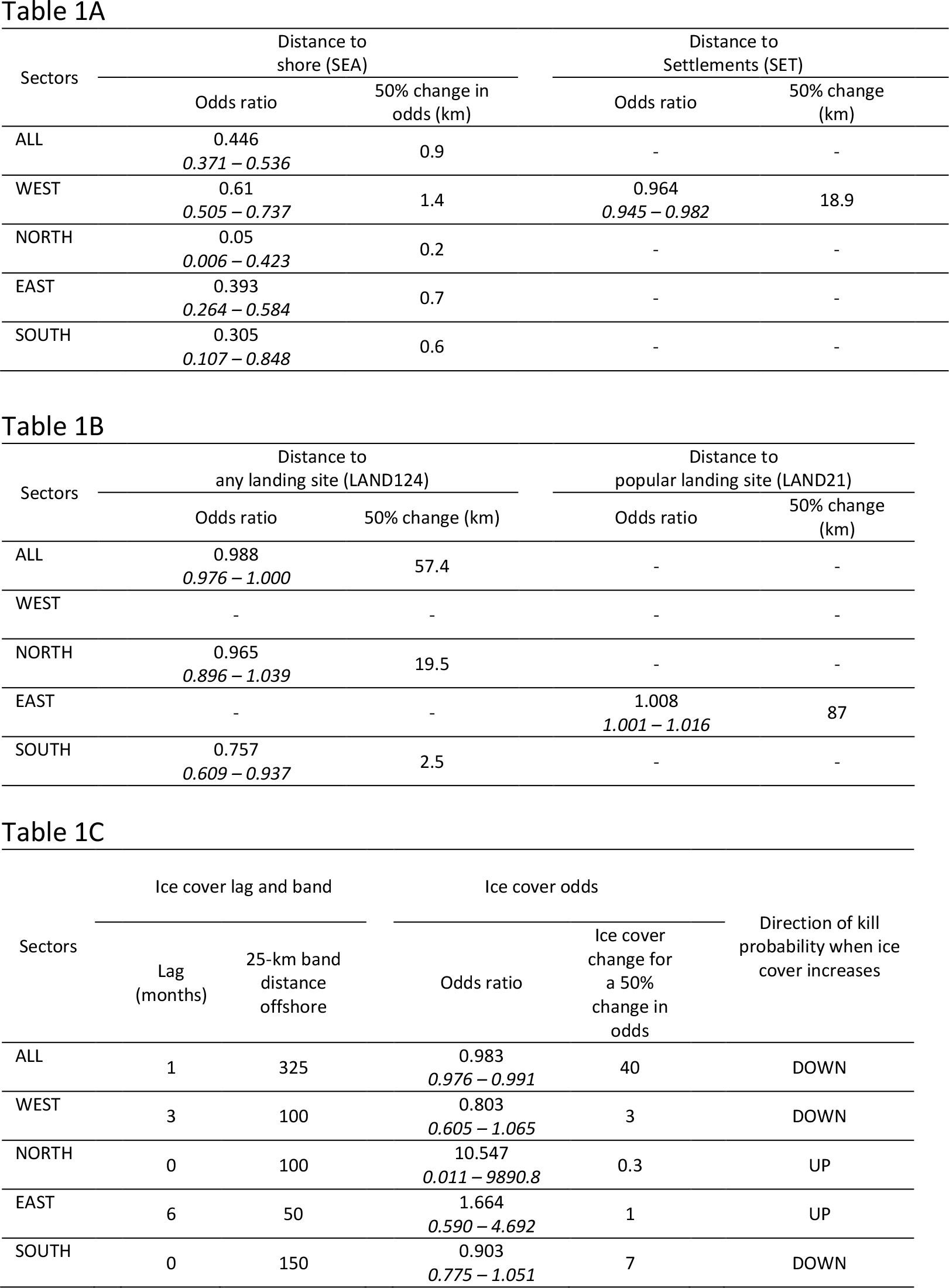
Relationships between distance covariates and ice cover with the odds ratios of a polar bear kill form the period 1987-2016. Tables 1A and 1B shows odds ratio, with 95% confidence intervals, per covariate unit (km) and the distances you have to move away from shore line, landing site, or settlement to reduce the odds for a kill with 50% (when odds ratio are larger than 1, this is not calculated as odds ratios increase). In Table 1C we have calculated and presented the drop in ice cover in the given ring and lag that will reduce the odds for a kill by 50%. Only results from the top model in each sector are shown. Results for all models see Table S5.

All four selected sector models, including the model for the pooled data set Sector All, identify distance to shore as an important predictor, i.e. odds ratio < 1 and confidence intervals for the variable SEA excluded the value 1. Odds dropped by 50% in the North sector when a theoretical bear moved only 200 m from the shoreline, 600 m in the East sector, 700 m in the East sector, 1.4 km in the West sector, and 900 m when all sectors were combined. These 50% reduction distances indicate that most kills occurred near shore and that proximity to shore was relatively hazardous to bears.

In the West sector the human-distance covariate SET, distance to human settlements, have odds ratio less than one, with a corresponding 95% confidence interval excluding one. The selected West model with its odds ratio of 0.964 [95% CI: (0.945, 0.982)], means that the odds of a bear kill decrease by 50% when the theoretical bear moved 18.9 km away from a settlement.

The other distance covariates lacked significant influence on the odds of a kill, except for distance from landing sites in the South sector, where an increase in distance of 2.5 km from any landing site (LAND124) resulted in a 50% drop in odds of a kill. For all sectors combined the required distance removed from a landing site for a 50% drop in odds ratio was 57.4 km.

For the four sectors separately, we found that odds of bear kills were influenced by ice cover in bands close to the shore. Namely, the best fitting discrete choice model contained the 100 km band in the West and North sectors, the 50 km band in the East sector, and the 150 km band in the South sector. In the West and the South an increase in ice cover of three and seven percent, respectively, in these bands resulted in a 50% decrease in the probability of a kill.

When considering ice time-lags, intended to evaluate the significance of prior ice concentrations on prediction of kills, increased ice cover three months prior to the kill reduced the probability of a kill in the West sector by approximately 20%. In the South sector, increased current ice cover (lag 0) reduced odds of a kill by approximately 10%.

In contrast, for the North and East sectors, the odds of a kill increased when ice cover increased 6 months and 0 months, respectively, prior to the kill. However, confidence intervals for these odds ratios are wide for some sectors, emphasizing the lack of reliable evidence across sectors (Table S5). When sectors are combined, a 40% increase in ice cover in the 25-km bands off the coast in the same month or one month prior to a kill resulted in a 50% drop in odds of a kill.

There were no significant effects of distance from trappers’ cabins on the odds ratio for a kill in any sector, or for all sectors combined.

## DISCUSSION

Availability of sea ice habitat around Svalbard declined greatly through the years represented in our study, resulting in more polar bears spending more time on land where they are largely food deprived (Stirling et al., 1999; Towns et al., 2009). Simultaneously, numbers of visitors to Svalbard increased dramatically (Norwegian Polar Institute, 2022). More bears on land for longer periods during which more people were accessing the same habitats could have been expected to increase the number of bear-human interactions, and the number of bears killed in defence of life and property. Despite a positive relationship between number of tourists and number of kills at a given time, the total numbers of bears killed did not increase over the years of the study and per-capita kills strongly declined. Whether by serendipity or because of conscious actions on the parts of managers and visitors, the temporal trend in kills is a surprising and hopeful outcome of this work.

This overall reduction in kills, despite greatly reduced sea ice habitat availability and more polar bears spending more time on land (Stirling et al., 1999; Hovelsrud et al., 2008; Towns et al., 2009), may reflect success of the Svalbard Environmental Act of 2001. This act prohibits people from “luring, pursuing or otherwise seeking out polar bears in such a way as to disturb them or expose either bears or humans to danger” (Norwegian Ministry of Climate and Environment, 2001). These requirements may make visitors more cautious and result in human behaviours that make lethal conflicts between humans and bears less likely.

There have been no studies focusing on human-bear interactions in Svalbard since 1993 (Gjertz et al., 1993), however, evidence suggests records of killed bears are comprehensive. Information about non-lethal interactions between polar bears and humans, however, is lacking. Incident data in Svalbard have prior to recent years only been collected when bears have been killed due to self-defence or loss of human life. Hence, we are unable to evaluate trends in total numbers (combined lethal and non-lethal) of incidents. The absence of such information is clearly a shortcoming that limits the understanding of the variety of bear/human interactions. Increased international collaboration stemming from the development of a coordinated circumpolar action plan to conserve polar bears have increased focus on the necessity to collect information on all kinds of incidents, including non-lethal ones (Polar Bear Range States, 2015).

The number of people visiting Svalbard has increased dramatically, especially after 2011 (Norwegian Polar Institute, 2022). The positive relationship between people arriving in Longyearbyen, the main settlement and the location of the airport, and fatal conflicts anywhere on Svalbard, indicates that the number of people arriving in Longyearbyen is a valid proxy for the number of people travelling in the wild, and thereby exposing themselves for potential interactions with polar bears. Although the vast majority of visitors to Svalbard stay in Longyearbyen, or go cruising on a variety of vessels, many depart from Longyearbyen to go trekking and camping in more remote areas where bears can be encountered. Data are lacking, however, to identify trends in the number of people travelling in the wild.

All sectors show a maximum of fatal conflicts in summer (July and August). In addition, there are lesser peaks in spring in the West and South sectors, and in April also in the East sector. These spring peaks coincide with the period when light returns and when snowmobile tourism peaks, and popular snowmobile trips runs from the main settlement Longyearbyen in the West to East sector areas of Agardh and Storfjorden, where polar bear females hunt with their COYs (cubs of the year) after den emergence in early April.

Although our records include 81 kills of problem bears, Wilder et al. (2017) found only 10 incidents where polar bears actually attacked people. Attacks were defined as “intentional contact by a bear resulting in human injury”. This means that the rest of the fatal incidents in our data set resulted from contacts with bears that were perceived as dangerous to humans, but where there were no actual attacks. Meetings between bears and humans can be placed on a continuum from a fatal conflict where bears and/or human lives are lost, to observations of bears made from a distance, not perceived as dangerous by the humans observing the bears. Consistent data on non-lethal encounters between people and polar bears would be important to collect for future management of polar bears in areas where people move around. The real nature of a meeting with bears can be hard to assess for anyone. Whether a situation is a real conflict incident and thus problematic, or an incident with no conflict, can be very hard to judge, and outcomes of interactions will always be partly dependent on the level of knowledge and experience of the humans involved. It is therefore vital that people moving around in bear country are trained in how to perceive the difference, how to accurately classify encounters as an observation, an interaction or a conflict; and how to respond appropriately (Hopkins et al., 2010; Wilder et al., 2017). This requires recording all encounters to enumerate, classify, and understand them, and to learn how to avoid having them escalate to the fatal stage.

The consistent finding that the odds for a problem bear kill is by far greater along the shoreline in all sectors was expected. The large majority of people that travel in the Svalbard wilderness never travel far from shore, and the West sector is the most populated and most trafficked part of Svalbard. Tourist peak seasons are April, which is peak snow mobile season, and summer months July and August. All sectors show peaks in these two periods of the year, while the time and place for kills shows a bit more variability in the West and South sector. Much of this variability can be explained by the annual pattern of sea ice cover around Svalbard. The western side of Spitsbergen is the most accessible part of Svalbard due to Atlantic water masses leaving the western part ice free most of the year, thus spreading tourist traffic more evenly across seasons than is the case in the north and the east.

Landing sites for tourists only affected the odds of fatal polar bear encounters in the South sector, where the odds of a fatality increase when approaching landing sites. The landing sites in the South sector are all within the Hornsund fjord, an area where historically there have been numerous observations of bears and bear tracks. Regular and frequent polar bear observations have been recorded at a permanent Polish research station in Hornsund established in 1957. This may be explained by the fact that the drift ice originating from the Barents Sea in the east, which historically has surrounded the southern tip of Svalbard on an annual basis, most often did not move further north than Hornsund (Larsen, 1986). And, although polar bears are regularly observed all along the western coast, where there are many tourist landing sites, Hornsund is only a few kilometer from Hambergbukta and the eastern fjord of Storfjorden by land, an area close to core denning areas and an area where many of the so-called “near-shore” bears spend large parts of their lives (Mauritzen et al., 2002).

It seems fair to conclude that increasing distances (i.e., distance covariates) predict better outcomes for bears, demonstrating a relationship between human and bears – namely, that bears enjoy better odds of *not* experiencing a DLP event by avoiding humans.

The decreased odds of a kill following an increase in ice cover in the sea ice bands 100-125 km offshore in the prior 4-6 months in the West sector could be explained by fewer bears being forced to stay ashore during winter in the west, and consequently also in peak tourist seasons in spring and summer. However, in contrast to the West and South sectors, where odds ratios is <1 with increasing ice cover, odds ratios are >1 in the North and East sectors as ice cover increases. This is hard to explain, as is what seems to be an effect on odds of a kill of sea ice cover in the outermost sea ice bands, 3-400 km offshore, in the same or the one month prior to a kill, when looking at all sectors combined. It could well be an artefact of our small sample size and the idiosyncrasy of ice covariates values and kill locations.

However, there seems to be different ecological strategies among bears in and around Svalbard, with a local ecotype and a pelagic ecotype (Mauritzen et al., 2001; Aars et al., 2017), differences that possibly could be expressed in the form of different migrational patterns as a response to variations in ice cover.

## CONCLUSION

Contrary to what we might of have expected, this study has revealed a favourable trend in incidental kills of polar bears in Svalbard. Our study also shows that kills have been most common in near shore areas, and that some areas are likely to see higher incident rates than others. Core denning areas, which would be areas of expected high density use, are moving north and east due to a change in the timing of sea ice arrival (Merkel and Aars, 2022). Areas where we have seen a historic high density of polar bear use, e.g., tracks in the Hornsund fjord area, might still be areas where we should expect a high number of interactions. Special attention may be required to protect bears in higher density use areas.

We emphasize that collection of data on all bear-human interactions, non-fatal interactions included, along with special focus on areas known to show higher polar bear use, will be important in order to give us a more complete understanding of the influence sea ice distribution may have on polar bear distribution. Systematic data collection of all bear-human interactions would increase sample sizes and may help explain which ice characteristics may forewarn of conditions that could lead to increased interactions. There is undoubtedly a link between the patterns of sea ice decline from areas near Svalbard and the numbers of bears people are likely to see on land. Kill data alone, however, were insufficient to resolve this relationship. Hence, continued monitoring of sector by sector ice patterns will be an important part of future monitoring efforts.

## ACKNOWLEDGMENTS

This study was funded by Svalbard Environmental Fund, the Norwegian Polar Institute, Polar Bears International, World Wildlife Fund, and San Diego Zoo. Thanks to James Wilder and Margrete Keyser for their help in validating records of problem bears, and to Harvey Goodwin and Max Kiönig at the Norwegian Polar Institute for producing shapefiles for ice rings and sectors, and for calculating monthly averages of sea ice concentrations.

## Notes

### Competing Interest Statement

The authors have declared no competing interest.

