## Supplemental figures and tables for "Relating Polar Bears Killed, Human Presence, and Ice Conditions in Svalbard 1987 – 2019"

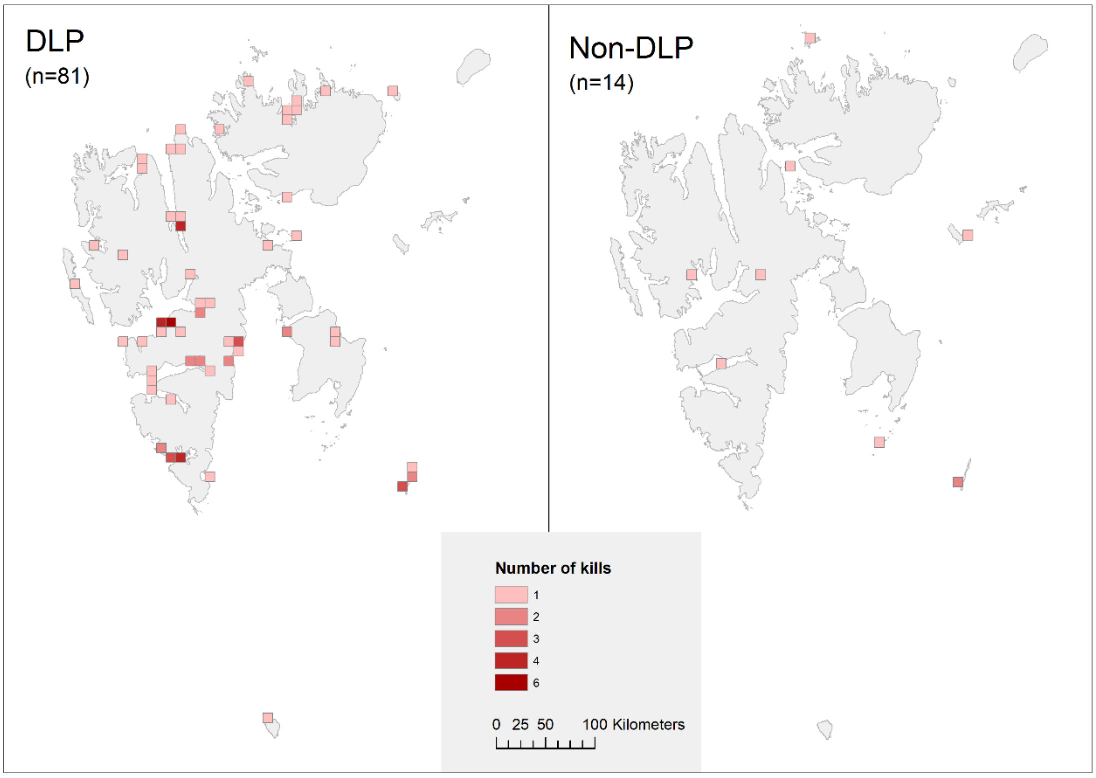


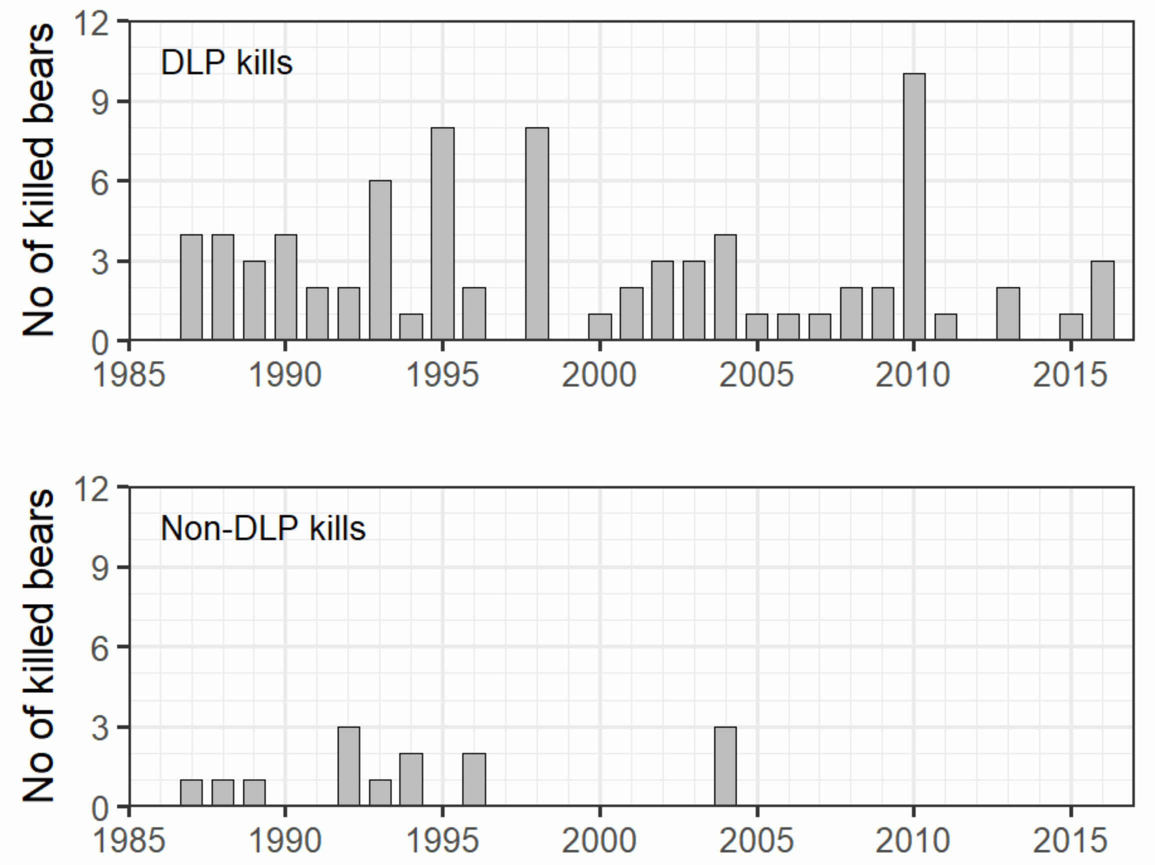


Figure S1 Kills of polar bears in Svalbard 1987-2016. Maps show locations aggregated over a 10 $km^2$ grid, and bar graphs show year of DLP and non-DLP kills.

Table S2 Existing data on tourism traffic at Svalbard in the period after 1995.

| Type of data | Coverage | Resolution | Source |
|---|---|---|---|
| People arriving in Longyearbyen | 1995 – present | Monthly | Visit Svalbard[1] |
| Guest nights in Longyearbyen | 1995 – present | Monthly | Visit Svalbard |
| Number of people disembarking from cruise ships | 1996 – present | Yearly | MOSJ[2] |
| Number of coastal landing sites for tourists | 1996 – present | Yearly | MOSJ |
| Number of people visiting areas outside management area 10[3] | 1998 – present | Yearly | MOSJ |
| Planned travel outside management area 10 | 1996 – 2017 | All planned trips | Governor of Svalbard |

[1] Main tourist operator in Longyearbyen, Svalbard

[2] Monitoring system for Svalbard and Jan Mayen (http://mosj.no/en/)

[3] Svalbard was until 2018 divided in 10 areas used to facilitate management actions

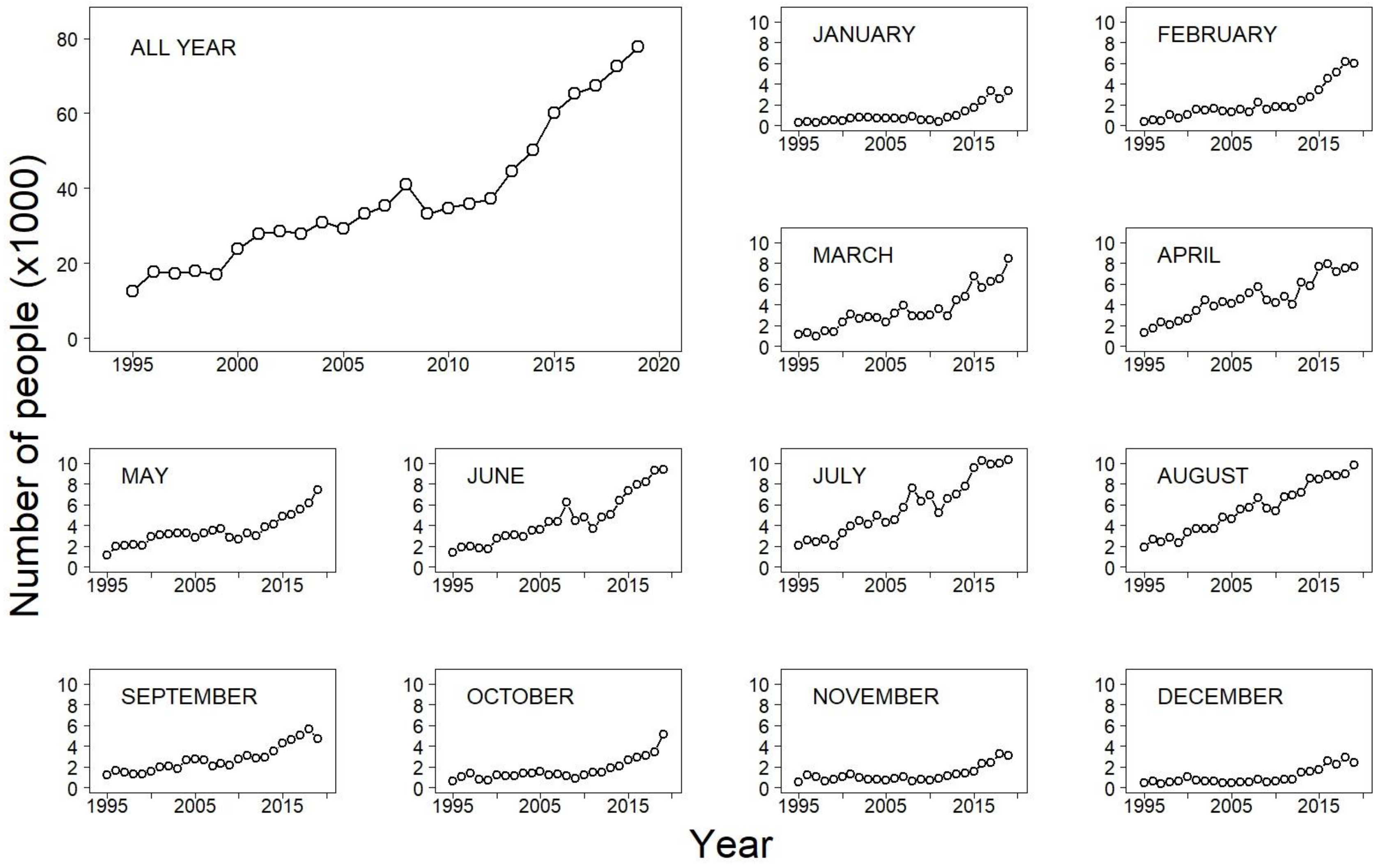


Figure S3 Number of people arriving in Longyearbyen per year (large panel) and each month (smaller panels) since 1995 (data from Visit Svalbard - https://www visitsvalbard.com).

Table S4 Counts of all kills from data set ALL and random points by sector, August 1987 – August 2016

| Kills | | Random points | | | | |
|---|---|---|---|---|---|---|
| Sector | Kills (ALL) | W | S | E | N | Total |
| W | 38 | 660 | 103 | 743 | 394 | 1,900 |
| S | 16 | 269 | 43 | 331 | 157 | 800 |
| E | 27 | 471 | 61 | 553 | 265 | 1,350 |
| N | 14 | 226 | 32 | 290 | 152 | 700 |
| Total | 95 | 1,626 | 239 | 1,917 | 968 | 4,750 |

Table S5 Odds ratios and confidence intervals of the odds ratios shown in Table 1.

| Sector | AIC Weights | Odds ratios for «Distance to» covariates | | | | Odds ratios for Ice Cover | | |
|---|---|---|---|---|---|---|---|---|
| | | Distance to shore (SEA) | Distance to Settlements (SET) | Distance to any tourist landing (LAND124) | Distance to popular tourist landing (LAND21) | Ice cover lag (months) | Ice cover distance offshore (km, outer band bin) | Kill odds ratio for 1% increase in ice cover |
| All | 0.165 | **0.446**<br>*0.371 - 0.536* | - | **0.988**<br>*0.976 - 1.000* | - | 1 | 325 | **0.983**<br>*0.976 - 0.991* |
| | 0.160 | **0.446**<br>*0.371 - 0.536* | - | **0.987**<br>*0.975 - 1.000* | - | 1 | 375 | **0.983**<br>*0.975 - 0.991* |
| | 0.126 | **0.447**<br>*0.372 - 0.537* | - | **0.987**<br>*0.975 - 0.999* | - | 0 | 375 | **0.983**<br>*0.976 - 0.991* |
| | 0.122 | **0.446**<br>*0.371 - 0.536* | - | **0.987**<br>*0.974 - 0.999* | - | 1 | 400 | **0.983**<br>*0.976 - 0.991* |
| | 0.116 | **0.445**<br>*0.370 - 0.535* | - | **0.989**<br>*0.977 - 1.001* | - | 1 | 350 | **0.984**<br>*0.977 - 0.992* |
| | 0.107 | **0.445**<br>*0.370 - 0.535* | - | **0.989**<br>*0.977 - 1.002* | - | 1 | 275 | **0.984**<br>*0.977 - 0.992* |
| | 0.101 | **0.446**<br>*0.372 - 0.537* | - | **0.986**<br>*0.974 - 0.998* | - | 0 | 400 | **0.984**<br>*0.976 - 0.991* |
| | 0.101 | **0.446**<br>*0.371 - 0.536* | - | **0.987**<br>*0.975 - 1.000* | - | 0 | 325 | **0.984**<br>*0.977 - 0.992* |
| W | 0.168 | **0.610**<br>*0.505 - 0.737* | **0.964**<br>*0.945 - 0.982* | - | - | 3 | 100 | **0.803**<br>*0.605 - 1.065* |
| | 0.130 | **0.608**<br>*0.503 - 0.735* | **0.962**<br>*0.944 - 0.980* | - | - | 6 | 125 | **0.802**<br>*0.602 - 1.069* |
| | 0.130 | **0.615**<br>*0.509 - 0.744* | **0.964**<br>*0.946 - 0.982* | - | - | 4 | 125 | **0.843**<br>*0.669 - 1.062* |
| | 0.127 | **0.608**<br>*0.502 - 0.736* | **0.964**<br>*0.945 - 0.982* | - | - | 5 | 100 | **0.855**<br>*0.707 - 1.034* |
| | 0.121 | **0.615**<br>*0.509 - 0.742* | **0.963**<br>*0.945 - 0.982* | - | - | 5 | 125 | **0.857**<br>*0.713 - 1.028* |
| | 0.116 | **0.616**<br>*0.511 - 0.743* | **0.964**<br>0.945 - 0.983 | - | - | 6 | 100 | **0.745**<br>*0.438 - 1.268* |
| | 0.106 | **0.619**<br>*0.513 - 0.745* | **0.964**<br>0.945 - 0.982 | - | - | 1 | 100 | **0.819**<br>*0.632 - 1.061* |
| | 0.103 | **0.608**<br>*0.502 - 0.736* | **0.964**<br>0.946 - 0.982 | - | - | 4 | 100 | **0.877**<br>*0.735 - 1.046* |
| N | 0.390 | **0.050**<br>*0.006 - 0.423* | - | - | - | 0 | 100 | **10.547**<br>*0.011 - 9890.8* |
| | 0.369 | **0.053**<br>*0.006 - 0.475* | - | - | - | 8 | 150 | **1.498**<br>*0.708 - 3.172* |
| | 0.240 | **0.072**<br>*0.009 - 0.593* | - | **0.965**<br>*0.896 - 1.039* | - | 8 | 150 | **1.525**<br>*0.333 - 6.992* |
| E | 1 | **0.393**<br>*0.264 - 0.584* | - | - | **1.008**<br>*1.001 - 1.016* | 6 | 50 | **1.664**<br>*0.590 - 4.692* |
| S | 0.585 | **0.301**<br>*0.107 - 0.848* | - | **0.757**<br>*0.609 - 0.940* | - | 0 | 150 | **0.903**<br>*0.775 - 1.051* |
| | 0.415 | **0.310**<br>*0.116 - 0.826* | - | **0.764**<br>*0.623 - 0.937* | - | 2 | 150 | **0.936**<br>*0.866 - 1.013* |
